# Robust transcriptional profiling and identification of differentially expressed genes with low input RNA sequencing of adult hippocampal neural stem and progenitor populations

**DOI:** 10.1101/2021.11.07.467608

**Authors:** Jiyeon K. Denninger, Logan A. Walker, Xi Chen, Altan Turkoglu, Alex Pan, Zoe Tapp, Sakthi Senthilvelan, Raina Rindani, Olga Kokiko-Cochran, Ralf Bundschuh, Pearlly Yan, Elizabeth D. Kirby

## Abstract

Multipotent neural stem cells (NSCs) are found in several isolated niches of the adult mammalian brain where they have unique potential to assist in tissue repair. Modern transcriptomics offer high-throughput methods for identifying disease or injury associated gene expression signatures in endogenous adult NSCs, but they require adaptation to accommodate the rarity of NSCs. Bulk RNA sequencing (RNAseq) of NSCs requires pooling several mice, which impedes application to labor-intensive injury models. Alternatively, single cell RNAseq can profile hundreds to thousands of cells from a single mouse and is increasingly used to study NSCs. The consequences of the low RNA input from a single NSC on downstream identification of differentially expressed genes (DEGs) remains largely unexplored. Here, to clarify the role that low RNA input plays in NSC DEG identification, we directly compared DEGs in an oxidative stress model of cultured NSCs by bulk and single cell sequencing. While both methods yielded DEGs that were replicable, single cell sequencing DEGs derived from genes with higher relative transcript counts compared to all detected genes and exhibited smaller fold changes than DEGs identified by bulk RNAseq. The loss of high fold-change DEGs in the single cell platform presents an important limitation for identifying disease-relevant genes. To facilitate identification of such genes, we determined an RNA-input threshold that enables transcriptional profiling of NSCs comparable to standard bulk sequencing and used it to establish a workflow for in vivo profiling of endogenous NSCs. We then applied this workflow to identify DEGs after lateral fluid percussion injury, a labor-intensive animal model of traumatic brain injury. Our work suggests that single cell RNA sequencing may underestimate the diversity of pathologic DEGs but population level transcriptomic analysis can be adapted to capture more of these DEGs with similar efficacy and diversity as standard bulk sequencing. Together, our data and workflow will be useful for investigators interested in understanding and manipulating adult hippocampal NSC responses to various stimuli.

## Introduction

The subgranular zone (SGZ) of the hippocampal dentate gyrus (DG) is a unique neurogenic niche in the adult mammalian brain (Denoth-Lippuner and Jessberger, 2021; Vicidomini et al., 2020). Neural stem cells (NSCs) in the SGZ give rise to functional new neurons throughout adulthood that contribute to hippocampal memory and affect regulation and could also be a source for endogenous tissue repair after injury or disease (McAvoy and Sahay, 2017; Miller and Sahay, 2019). Over the past decade, transcriptional analysis using high-throughput RNA sequencing (RNAseq) technology has dramatically expanded knowledge of NSC molecular characteristics. For example, studies using prospectively identified stem and progenitor populations have uncovered previously unknown cell lineage relationships (Baser et al., 2019; Berg et al., 2019; Dulken et al., 2017; Llorens-Bobadilla et al., 2015). Other studies using more unbiased approaches have revealed regional and even cell-specific transcriptional differences or transcriptional changes during development, adult neurogenesis, or aging (Artegiani et al., 2017; Dulken et al., 2019; Hochgerner et al., 2018; Mizrak et al., 2019; Shin et al., 2015; Yuzwa et al., 2017; Zywitza et al., 2018). As studies of the NSC transcriptome expand, researchers are faced with an increasing variety of options for how to accomplish transcriptional profiling of this small, but critical, cell population.

Current major challenges to transcriptional profiling of NSCs include their relative sparsity in vivo and their transcriptional similarity to astrocytes. Both of these challenges have made bulk RNAseq of prospectively isolated NSCs a less attractive approach as it requires large cell number input and prospective isolation of the desired population. Instead, single cell RNAseq (scRNAseq) has emerged as the preferred technique to begin overcoming the above barriers. This approach uses the very small amounts of RNA present in single cells to generate thousands of individual cell transcriptomes with massively paralleled sequencing. scRNAseq studies of the adult mouse SVZ and SGZ have identified rare subpopulations of cells, as well as dynamic changes in gene expression at different developmental stages, maturation states, and regional locations (Artegiani et al., 2017; Llorens-Bobadilla et al., 2015; Shin et al., 2015). Several studies have adapted analytical methods from standard bulk sequencing to accommodate the technical challenges presented by using such low input to profile cells from other lineages (Law et al., 2014; Love et al., 2014; Robinson and Oshlack, 2010). In addition, many pioneering studies in other cell types have also developed novel processing and analysis tools specifically designed to facilitate detection of cell heterogeneity or chronological mapping of developmental trajectories using scRNAseq transcriptomes (Finak et al., 2015; Qiu et al., 2017; Shin et al., 2015; Trapnell et al., 2013).

While these recent studies show that scRNAseq is a powerful approach to characterize differences between individual cells, it is not yet clear how effective it is for uncovering population-level changes in gene expression. Identification of differentially expressed genes (DEGs) induced by variables like injury or gene expression manipulation is critical to understanding the mechanisms underlying NSC function in both disease models and in healthy brains. It seems logical that the low input of scRNAseq would affect DEG discovery compared to standard bulk RNAseq, where DEG analysis in transcriptomics was first developed (Arzalluz-Luque et al., 2017; Bhargava et al., 2015). However, beyond a few recent studies directly comparing the efficacy of DEG identification by various statistical methods on single cell datasets, there is little information available directly comparing the quality of DEG analysis in scRNAseq versus bulk RNAseq (Mou et al., 2020; Wang et al., 2019; Ziegenhain et al., 2017). Here, we directly compare scRNAseq with bulk level RNAseq of cultured NSCs in a model of oxidative stress in vitro to evaluate DEG identification across sequencing approaches. We surprisingly found little overlap in DEGs identified by scRNAseq and bulk RNAseq, despite using the same source samples. While subsequent experiments showed that DEGs from both approaches were replicable, we found that scRNAseq identified DEGs among genes that show a more moderate fold change and high relative transcript count when compared to the bulk RNAseq approach. Because many studies of DEGs would specifically benefit from identification of higher fold change transcripts which are more moderately expressed, we adapted and validated a limiting cell (lc) RNAseq approach for sequencing DG NSCs isolated from individual adult mice with similar reliability as more bulk RNAseq-like approaches. We further demonstrate the utility of this method by applying it to transcriptome profiling of NSCs and their intermediate progenitor cell (IPC) progeny from single adult mouse hippocampi after a lateral fluid percussion injury (LFPI) model of traumatic brain injury (TBI).

## Results

### DEGs identified by scRNAseq versus bulk population level RNAseq are different but accurate

To compare detection of differential gene expression by scRNAseq and bulk RNAseq of cultured cells, we used an in vitro model of oxidative stress with adult mouse DG-derived NSCs. We treated cultured NSCs derived from male and female adult mouse DGs with H_2_O_2_ to induce oxidative stress or with vehicle and then harvested cell pellets from 3 biological replicates (1 male, 2 female) per treatment. Harvested cells were then subdivided into two processing streams: RNA extraction for bulk RNAseq of pooled cells or direct RNAseq of individual cells (scRNAseq) using a 10X Chromium platform (**Figure 1**).

**FIGURE 1.**
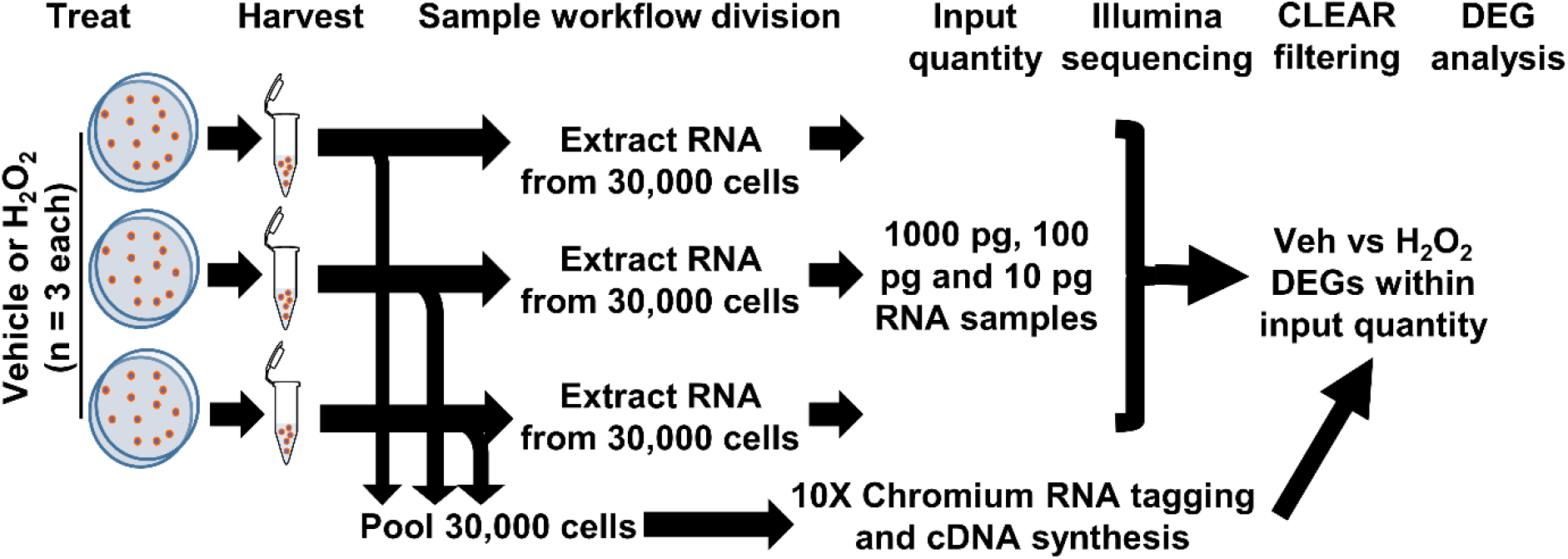
Workflow for cultured NSCs in scRNAseq and bulk RNAseq. NSCs were treated with H_2_O_2_ or vehicle in triplicate. 10,000 cells from each biologic replicate were pooled for a total of 30,000 cells applied to the 10X Chromium scRNAseq platform and subsequent DEG analysis. RNA was extracted from 30,000 cells of each biologic replicate and 10 pg, 100 pg, and 1000 pg (1 ng) were used in RNAseq with subsequent CLEAR filtering and DEG analysis.

In scRNAseq, a total of 26,916 cells were captured and sequenced. UMAP analysis of both H_2_O_2_-treated and control samples revealed 10 different subpopulations characterized by gene expression profiles linked to GO terms consistent with specific stages of the cell cycle, phases of quiescence, differentiation, or response to injury (**Figure 2A–B** and **Supplementary Table 1**). The majority of cells were in G_1_ (22%), G_2_/M (21%), or S (6.5%) phase of the cell cycle. 17.5% of cells were in one of two detected quiescent phases (G_0d_ and G_0r_). 16% of the cells appeared to be in an intermediate state that we characterize as transitioning to/from the cell cycle (T) and the remaining 17% of cells were differentiating (3%, D1 and D2) or responding to injury (14%, I and A). Cells from both treatment groups were present in all clusters (**Figure 2A**). However, vehicle treated cells were mostly concentrated in the cycling clusters while H_2_O_2_ treatment resulted in a notable shift away from cycling clusters to quiescent and apoptotic/injured clusters.

**FIGURE 2.**
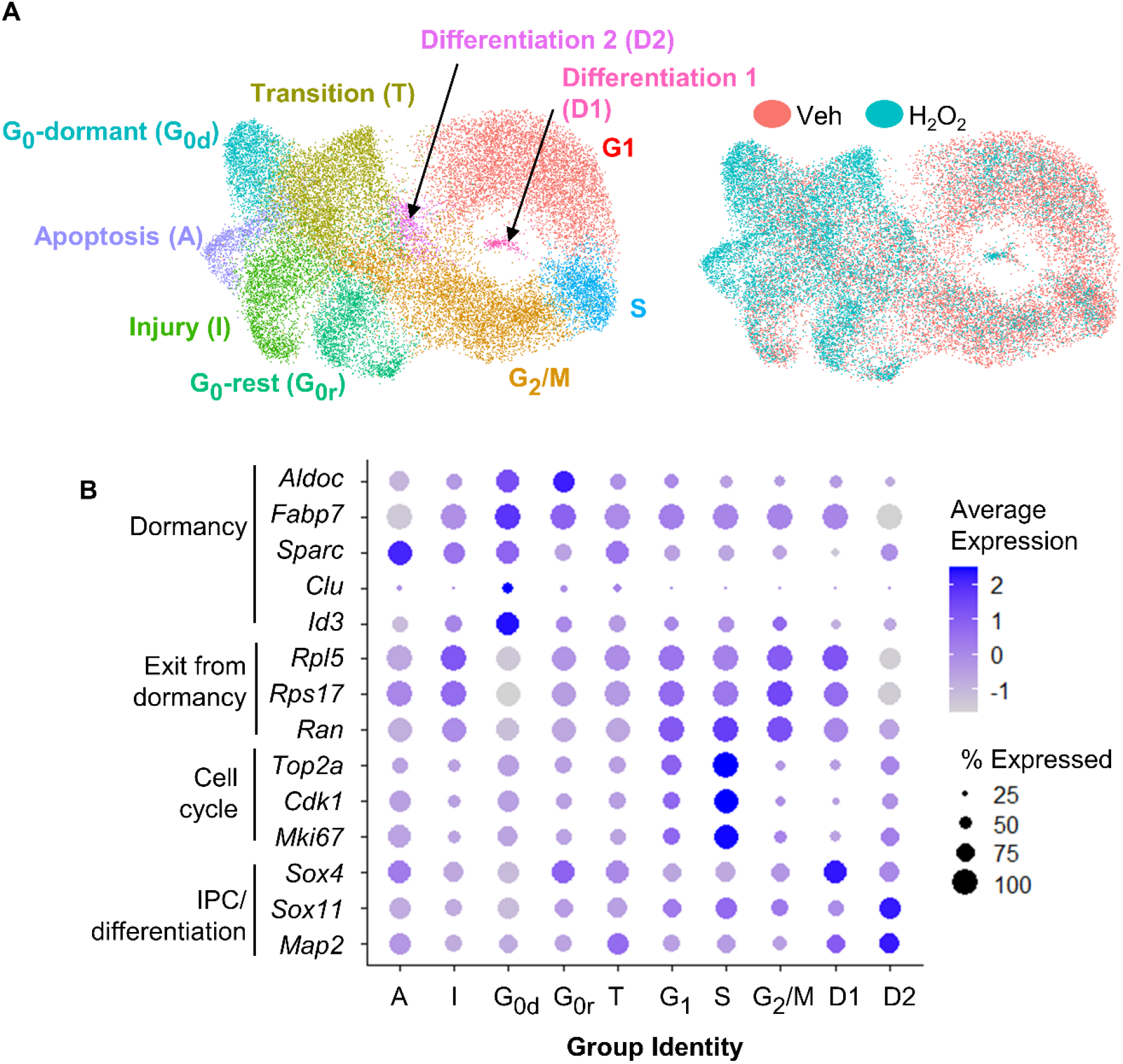
scRNAseq of cultured NSCs in an in vitro model of oxidative stress. **(A)** UMAP-of vehicle and H_2_O_2_-treated cultured NSCs yielded 10 subpopulations defined by gene expression profiles consistent with GO terms associated with various stages of the cell cycle, levels of quiescence or response to injury (left). UMAP comparison of H_2_0_2_-treated versus vehicle-treated NSCs indicated a shift towards quiescence, apoptosis, senescence, and injury response following oxidative stress (right). **(B)** Dot piot visualization of average expression of and percent of cells expressing select genes known to be associated with Quiescence versus activation further confirmed subpopulation identities.

To confirm the cluster identities of the dormant and cycling cells, we looked at expression of known markers of dormancy and progression through the cell cycle. The largest subpopulation, which we designated as G_1_ based on GO analysis, was characterized by moderate to high expression of genes involved in exit from dormancy, such as *Rpl5, Ran* and *Rps17* (Harris et al., 2021), as well as moderate expression of cell cycle genes such as *Top2a, Cdk1* and *Mki67* (**Figure 2A–B** and **Supplementary Figure 1A–B**) (Diril et al., 2012; Nielsen et al., 2020; Sun and Kaufman, 2018). The S phase population was characterized by a distinct upregulation of cell cycle gene expression (**Figure 2B** and **Supplementary Figure 1B**). The G_2_/M cluster showed continued high expression of exit from dormancy genes coupled with sharp downregulation of cell cycle genes relative to G_1_ and S (**Figure 2A–B** and **Supplementary Figure 1A–B**). Two clusters of G_0_-like cells were found, both of which showed upregulation of genes linked with GO terms such as ion homeostasis and metabolic processes. The two clusters differed most notably in expression of genes associated with transition to/from deeper quiescence. Specifically, G_0_-dormant was characterized by high expression of quiescence-associated genes *Fabp7, Aldoc, Sparc, Clu* and *Id3* (Artegiani et al., 2017; Borrett et al., 2020; Dulken et al., 2017; Urbán et al., 2019), coupled with low expression of exit from dormancy-associated genes *Rpl5, Rps17* and *Ran* (**Figure 2A–B** and **Supplementary Figure 1A and C**) (Harris et al., 2021). G_0_-rest, in contrast, showed an upregulation of exit from dormancy genes and slight suppression of quiescence genes (**Figure 2A–B**). The transition group (T) was intermediate in expression of markers of dormancy and cell cycle, supporting its assignment as a transitional state between quiescent G_0_ states and the cell cycle (**Figure 2A–B**). The two clusters of differentiating cells (D1 and D2) showed upregulation of genes associated with progenitor cell differentiation (*Sox4, Sox11* and *Map2*) (**Figure 2A–B** and **Supplementary Figure 1D**) (Artegiani et al., 2017; Hochgerner et al., 2018; Shin et al., 2015). Last, both injured and apoptotic clusters showed expression of genes associated with cell injury and death processes such as *Srxn1* and *Phlda3* respectively (**Figure 2A–B** and **Supplementary Figure 1E and F**) (Bell et al., 2015; Kawase et al., 2009).

**Supplementary Figure 1.**
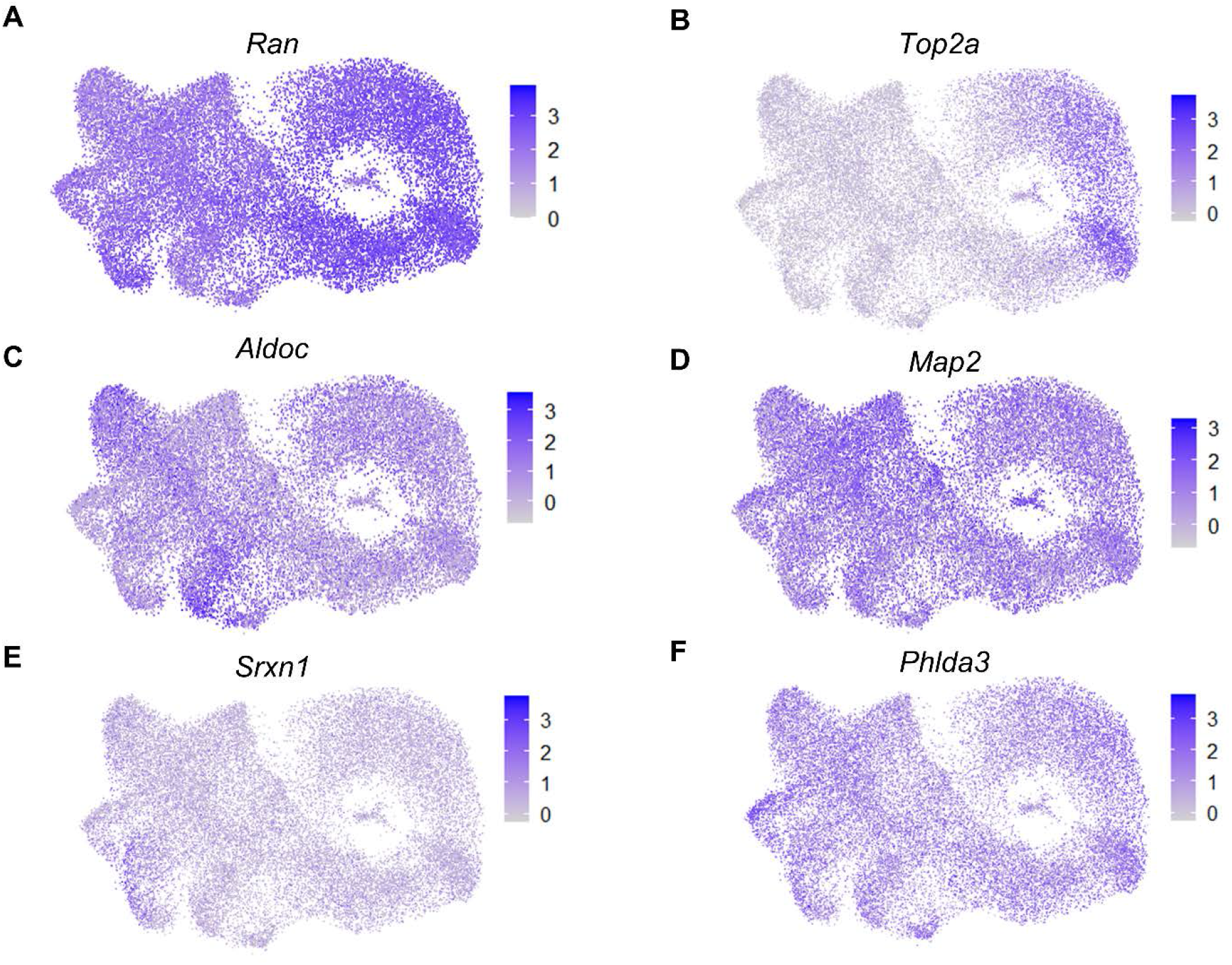
Feature plots of characteristic genes for subpopulations identified by scRNAseq of NSCs with and without H_2_O_2_-induced oxidative stress in vitro. **(A)** *Ran* as a representative gene for exit from dormancy **(B)** *Top2a* as a representative gene for cycling cells. **(C)** *Aldoc* as a representative gene for dormancy/quiescence. **(D)** *Map2* as a representative gene for differentiating cells. **(E)** *Srxn1* as a representative gene for injured cells. **(F)** *Phlda3* as a representative gene for apoptosis.

We next identified DEGs between H_2_O_2_ and vehicle treated cells using both the scRNAseq data and RNAseq from pooled cells using 1 ng total input RNA, an amount which yields sequencing data in the bulk RNA sequencing range (Walker et al., 2020). For scRNAseq, all clusters were combined within treatment for DEG analysis using the standard Wilcoxon test. 299 DEGs were identified between H_2_O_2_ and vehicle treated NSCs using scRNAseq (**Supplementary Table 2**). 1ng RNAseq data was pre-processed using the coverage-based limiting-cell experiment analysis (CLEAR) pipeline to eliminated unreliable, lowly expressed transcripts (Walker et al., 2020) then DEGs were identified using DESeq2. The 1ng RNAseq comparison yielded 790 DEGs between H_2_O_2_ and vehicle treated NSCs (**Figure 3A** and **Supplementary Table 2**). Comparison with the DEGs identified between sequencing methods revealed only 93 genes that were common to both platforms. Not surprisingly, most of the non-overlapping genes (697) were unique to the higher RNA input platform of 1 ng RNA, implying an expected greater sensitivity for DEG detection with greater RNA input. More unexpectedly, though, scRNAseq identified 206 unique DEGs compared to the 1ng input RNAseq. This low overlap between sequencing approaches from the same source NSCs implied either that one method was calling numerous false DEGs or that the two platforms had strongly different biases in what DEGs they can detect.

**FIGURE 3.**
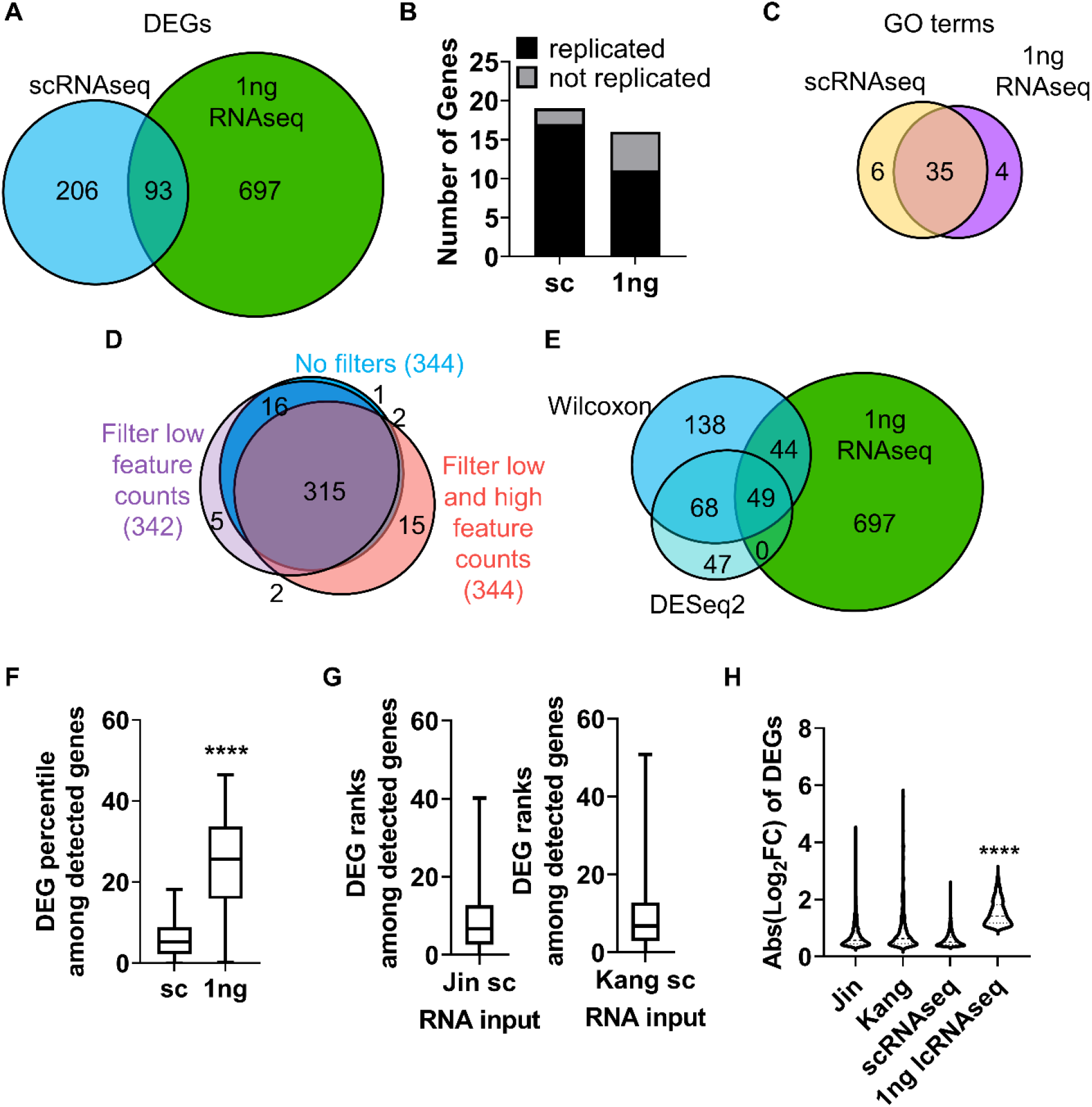
RNA input leads to bias in DEG discovery. **(A)** Venn diagram of D“’EGs identified by scRNAseq and 1 ng RNAseq with CLEAR filtering. **(B)** qRT-PCR analysis corroborated the majority of DEGs identified by scRNAseq and 1 ng RNAseq in cultured NSCs following H 0r induced oxidative stress. X^2^ contingency test (df = 1) = 2.331, p = 0.127. **(C)** Venn diagram of GO terms associated with DEGs identified by scRNAseq and 1 ng RNAseq. **(D)** Venn diagram of DEGs identified by scRNAseq with no filtering, filtering for low feature counts, and filtering for both low and high feature counts (i.e. the default for Seurat analysis). **(E)** Venn diagram of DEGs identified by scRNAseq using the Wilcoxon test or DESeq2 compared with DEGs identified by 1ng RNAseq. **(F)** DEGs were ranked by percentile of average transcript count level relative to all detected gene counts. The highest count genes are the lowest number percentile (i.e. highest count relative to all detected genes would be in the 1^st^ percentile). Box plot shows median and quartiles of percentiles of DEGs. DEGs identified by scRNAseq averaged the 6^th^ percentile while DEGs identified by 1ng RNAseq averaged the 25^th^ percentile. ****p<0.0001 unpaired t-test. **(G)** DEGs identified in scRNAseq data from Jin et al. {left) and Kang et al. (right) were ranked by transcript level percentile relative to all detected genes in those datasets. Similar to our scRNAseq data, the majority of DEGs were limited to the top 10% of expressed genes. **(H)** Comparison of fold changes in average transcript count between treatment groups for DEGs from all datasets indicated that DEGs identified by 1 ng RNAseq showed significantly larger fold changes in gene expression than DEGs identified by any of the scRNAseq datasets. ****p<0.0001 Dunnett’s multiple comparison test.

To determine how accurate and replicable the DEGs from scRNAseq and 1ng RNAseq were, we repeated H_2_O_2_/vehicle treatment of cultured NSCs in two more independent replications and analyzed gene expression of the top upregulated DEGs identified by scRNAseq and 1 ng RNAseq using quantitative real time PCR (qRT-PCR). Of the top 20 genes, we achieved effective primers for 19 of the top 20 scRNAseq DEGs and 16 of the 1 ng RNAseq DEGs. The majority of genes identified by both scRNAseq and 1 ng RNAseq were confirmed to be upregulated with qRT-PCR and there was no significant difference between methods in the number of genes replicated (**Figure 3B** and **Supplementary Figure 2A**). To further understand what distinguished DEGs called by scRNAseq versus 1ng RNASeq, we compared overlap of GO terms for upregulated genes in each platform. GO terms showed a high degree of overlap (35 out of 45 distinct categories) in the biological processes represented by the genes identified with both scRNAseq and 1 ng RNAseq (**Figure 3C** and **Supplementary Table 2**). This overlap in biological processes being triggered by oxidative stress, coupled with the high replicability of individual DEGs, implies that both platforms likely generated accurate DEGs that reflect true changes in cell activity. However, the low degree of overlap in the individual DEGs identified by the 2 platforms indicates that some form of bias affected the type of genes identified by each platform.

### RNA input amount drives bias in DEG discovery

The first possibility we considered for the difference in DEG identification between sequencing platforms was that it was an artefact of the different data analysis and statistical techniques used to identify DEGs in scRNAseq versus 1ng RNAseq data. scRNAseq analysis involves several layers of filtering to compensate for technical limitations of low capture efficiency, high dropouts, and doublets. Two commonly used quality controls (QCs) set minimum and maximum thresholds for Unique Molecular Identifiers (UMIs) and feature counts to filter out damaged cells or doublets (Stuart et al., 2019). However, such filtering relies on predetermined values and could overzealously filter out valid cells on the ends of the spectrum of true cellular differences in RNA content and complexity. To determine if QC limits altered DEG discovery, DEG analysis was performed on scRNAseq data with and without filtering for low (1000 counts/cell) and high (2500 counts/cell) feature counts (**Figure 3D**). Removing both high and low feature counts resulted in a total of 344 DEGs with 92% overlap compared to fully filtered data (**Figure 3D**). Filtering out cells with only low feature counts yielded 342 DEGs with almost 93% of those genes also identified when all QCs are employed (**Figure 3D**). This comparison revealed high overlap in the DEGs identified with and without the various filters, indicating that QC filtering of scRNAseq data was not introducing the divergence in DEG discovery between sequencing approaches.

We next considered whether the different statistical tests used in scRNAseq and 1ng RNAseq might have contributed to differential DEG identification. For scRNAseq data, we used the default in Seurat pipeline implementation of the Wilcoxon test. Bulk RNAseq, in contrast, is typically (and was in our case) analyzed using DESeq2. To determine if the choice of statistical test affected the DEGs identified, we compared DEGs called by DESeq2 in both scRNAseq and 1ng RNAseq. DEG discovery using DESeq2 of scRNAseq data resulted in far fewer genes (a little over half the number identified using the Wilcoxon test), but 71% of those genes were common to both statistical methods (**Figure 3E**). DESeq2 has been noted to be more restrictive (Mou et al., 2020) so the fewer DEGs from that analysis is not surprising. While the two different methods of statistical analysis of scRNAseq data yielded highly overlapping sets of genes, they both only marginally overlapped (about 30%) with DEGs identified with DESeq2 in the 1 ng RNAseq dataset (**Figure 3E**). This comparison suggests that use of Wilcoxon versus DESeq tests is not likely the source of DEG discordance between scRNAseq and 1ng RNAseq approaches. Rather, these data suggest that there is something inherently different in the data generated with scRNAseq versus 1 ng RNAseq that is leading to different biases in DEG discovery with each approach.

To further explore the differences in DEGs identified in scRNASeq versus 1ng RNAseq, we next looked at the relative transcript count level of DEGs. Gene expression level within a cell impacts the likelihood of transcript capture and is known to significantly influence DEG analysis in single cell studies (Mou et al., 2020). Indeed, the low RNA input of scRNAseq is expected to result in high drop out and a consequent overall reduction of detected genes and therefore DEGs. The identification of DEGs in scRNAseq that were not identified via 1ng RNAseq in our data, however, is more unexpected. To determine if gene expression level led to the discordance in DEG discovery here, the averaged transcript counts per cell of DEGs identified in each sequencing approach were compared to those for all detected genes within that same approach (**Supplementary Figure 2B and C**). Genes that were not expressed in any cells were excluded to limit bias from zero inflation, especially in scRNAseq. To enable direct comparison between these two datasets, we then converted the transcript counts of DEGs within each sequencing platform to percentiles relative to the average transcript counts of all detected genes. In this comparison, genes with the highest average transcript count per sample comprise the top percentiles (e.g. 1st %ile) of counts (**Figure 3F**). DEGs identified by 1 ng RNAseq had a higher median transcript count percentile value (~25%ile) and spanned a wider percentile range of transcript count levels than those identified by scRNAseq, which were limited to the top 5-10% of detected transcripts (**Figure 3F**). It is important to note that these comparisons were made after eliminating genes with no counts, showing that a strong bias for high count genes persists in scRNAseq data even when undetected genes are excluded. To determine how widespread this bias for high count genes might be, we analyzed two other, published scRNAseq datasets from different model systems. Kang et al. performed droplet-based scRNAseq of human peripheral blood mononuclear cells treated with vehicle or interferon beta ex vivo (Kang et al., 2018). Jin et al. performed scRNAseq of all CD45 highly positive cells isolated from young and aged mouse dentate gyrus (Jin et al., 2021). Despite using different species, source tissue and manipulations, we found that scRNAseq DEG discovery in these two studies was similarly limited to the top 5-10% of high count transcripts (**Figure 3G**).

**Supplementary Figure 2.**
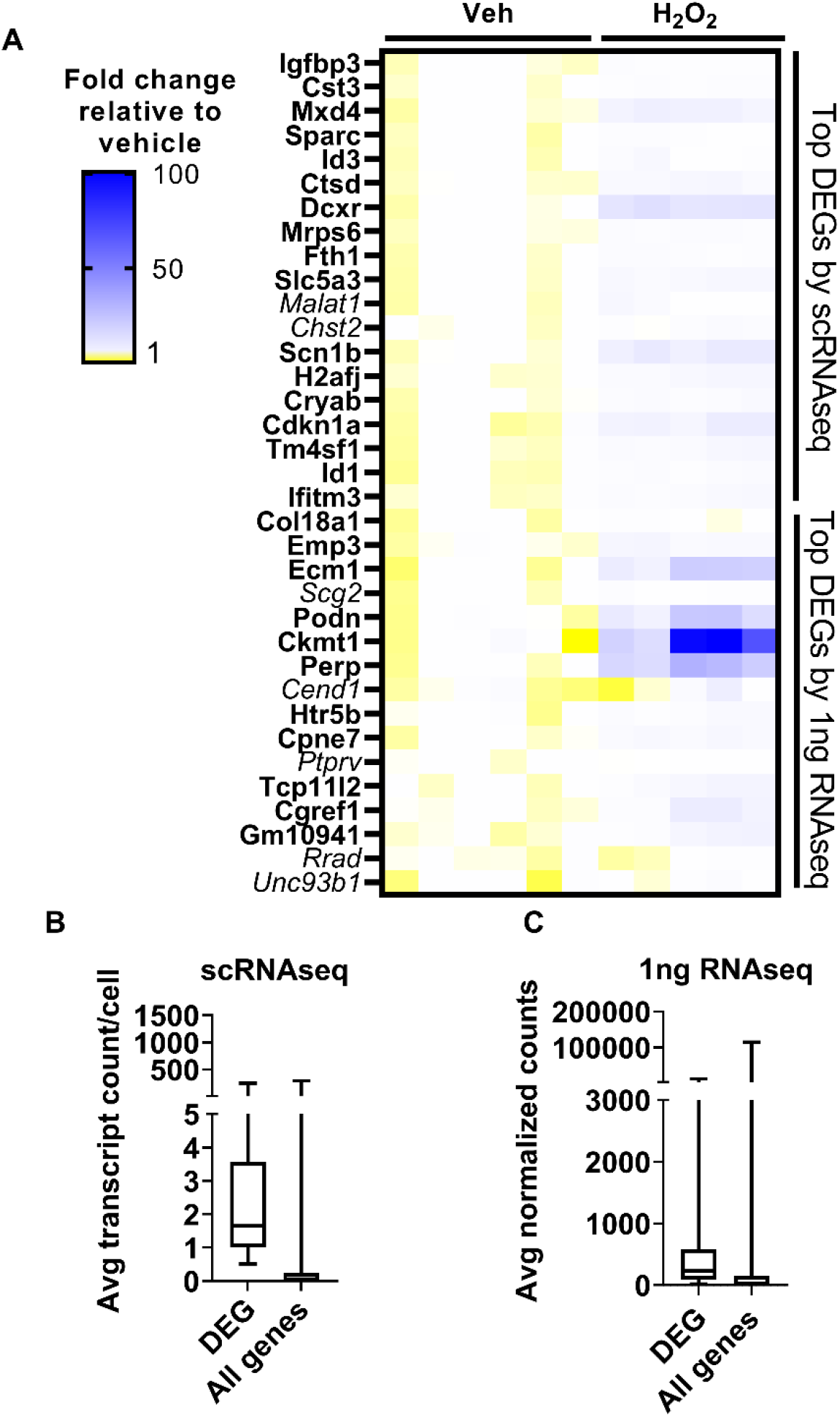
Gene expression levels impact DEG identification. **(A)** qRT-PCR of the top DEGs identified by scRNAseq or 1 ng RNAseq of cultured NSCs treated with H_2_O_2_ or vehicle show significant upregulation for most genes. Genes not significantly different between groups by Mann-Whitney test are italicized. N = 5-6 well replicates from 2 independent experiments. **(B)** Box plots show the median and quartiles of transcript count per cell for DEGs and all genes in scRNAseq data. **(C)** Box plots show the median and quartiles of transcript count per cell for DEGs and all genes in 1 ng RNAseq data. Genes with zero expression in all cells were eliminated for (B) and (C).

We next analyzed the relative fold change in counts of DEGs identified by scRNAseq and 1ng RNAseq. We found that scRNAseq in our experiment and in the two independent, unrelated comparison datasets yielded DEGs with lower fold changes than 1 ng RNAseq did. scRNAseq dataset DEGs showed an average fold change mostly between 1.5 and 2.5 (**Figure 3H**). Our 1 ng RNAseq data, in contrast, had a 3.0 fold change on average. Combined with the above findings on relative count level of DEGs, these findings suggest that the divergence in DEG detection between our scRNAseq and 1ng RNAseq data is driven by bias in scRNAseq data for detection of DEGs that derive from high count genes that show a more moderate fold change and in 1ng RNAseq data for DEGs from more moderate count genes that show a higher fold change between groups.

### RNAseq determination of optimal RNA input for DEG discovery

The comparison of scRNAseq and 1ng RNAseq shows that while both methods accurately identify DEGs, RNA input biased the types of DEGs that were detected. In studies seeking to identify specific causative genes implicated in a manipulation (e.g. disease model, drug treatment, altered gene expression), discovering genes that have high fold change and low expression level is particularly advantageous. Our data suggest that scRNAseq may be distinctly ill-suited for this purpose and a bulk RNAseq approach would be better. However, in the case of in vivo NSCs, the number of cells isolated from a single mouse is lower (100s-1000s) than that which is typically necessary for bulk RNAseq (100,000s). We used our in vitro oxidative stress model to determine the consequences of reduced RNA input, like that which may be encountered in a limiting cell (lc) RNAseq analysis of in vivo NSCs, for DEG detection. From the same samples, we compared transcriptional profiles derived from 1ng, 100pg and 10pg of RNA input. Similar to our analysis of the 1ng samples above, 100 and 10 pg RNAseq data was pre-processed using the CLEAR pipeline to eliminated unreliable, lowly expressed transcripts (Walker et al., 2020). Principal component analysis (PCA) showed that biological replicates completely overlapped with each other, indicating that the cultured NSCs were fairly uniform populations of cells, regardless of passage number or sex of the cells (**Figure 4A**). There was a clear separation between H_2_O_2_- and vehicle-treated NSCs on principal component (PC) 1 (**Figure 4B**). The clearest separation, however, was by RNA input level, along PC2 (**Figure 4C**).

**FIGURE 4.**
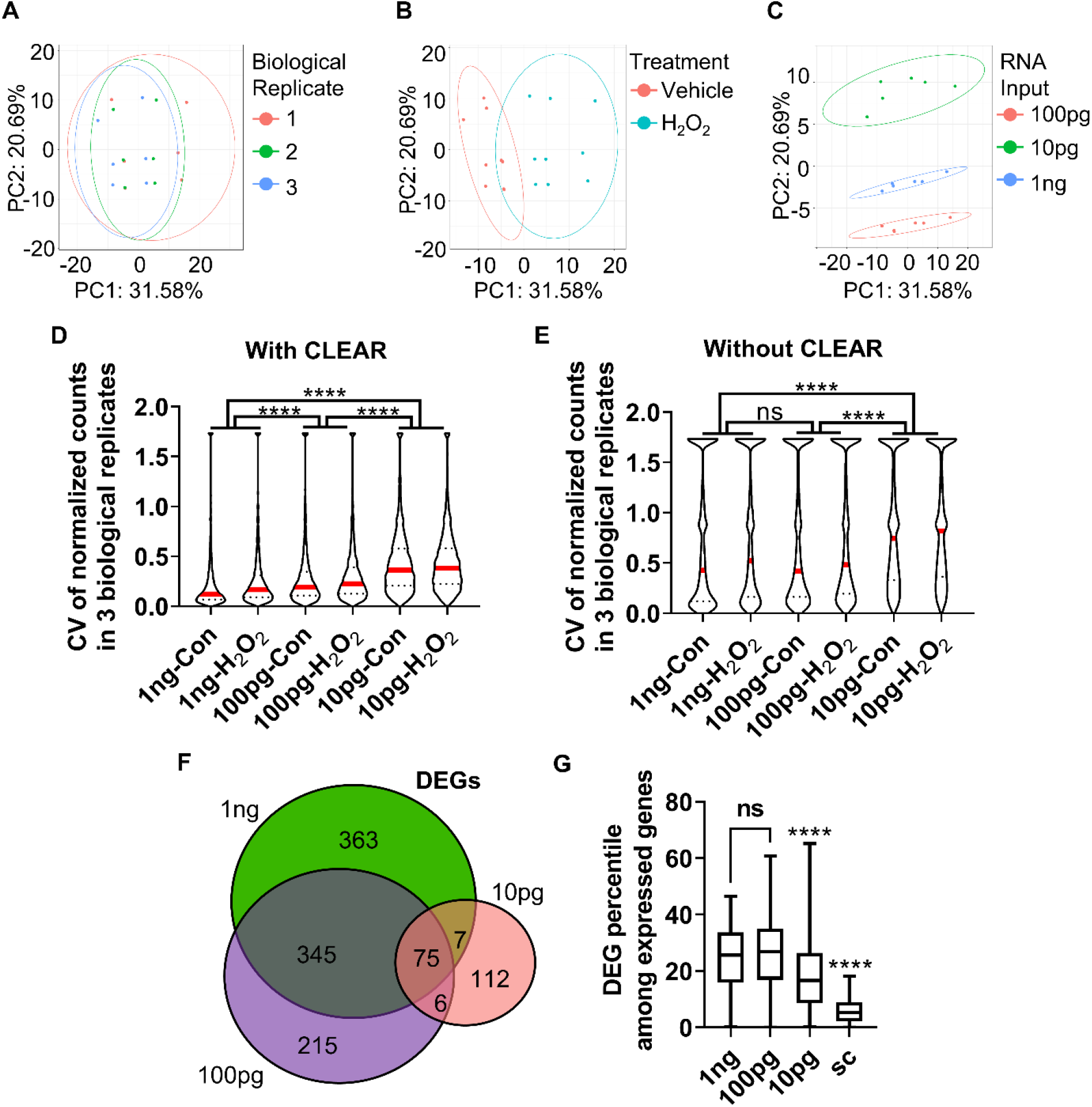
Determination of optimal input RNA amount with RNAseq and CLEAR that preserves unbiased DEG identification. **(A)** Biological replicates of cultured NSCs overlap in PCA. **(B)** PCA of cultured NSCs shows that cells undergoing oxidative stress diverge from control cells along principal component (PC) 1. **(C)** PCA of samples based on RNA input amount leads to significant separation between samples along PC2. **(D)** CVs were inversely related to RNA input amount. The difference in median CV between all RNA input levels were statistically significant. Median (red line) and quartiles (dotted line)) are shown within violin plots of CV for all genes detected in all samples. **(E)** Analysis of the CV with vehicle and H_2_O_2_-treated samples of all three RNA inputs before CLEAR filtering shows substantially greater CVs at all RNA input levels, confirming the utility of CLEAR preprocessing. Median (red line) and quartiles (dotted line) are shown within violin plots of CV for all genes detected in all samples. **(F)** Venn diagram of DEGs identified in all three RNA input levels show majority overlap between 1 ng and 1 00pg but low overlap with 10pg. **(G)** Comparison of transcript count percentiles for DEGs identified at the three different RNA input levels show no significant difference in DEG percentile between 1 ng and 1 00pg RNA inputs. ****p<0.0001 One-Way ANOVA with Kruskal-Wallis test and Dunn’s multiple comparisons in (D), (E), and (G).

Comparison of the median coefficient of variation (CV) between biological replicates showed that CV increased significantly with decreasing RNA input (**Figure 4D**). However, the difference in median CV was much more moderate between 1 ng and 100 pg (~46% increase from 1ng to 100pg) versus 1 ng and 10 pg (~77% increase). These data indicate lower precision and reproducibility as RNA input is reduced, particularly when it drops below 100 pg (**Figure 4D**). Notably, CV was substantially larger in all conditions when data were analyzed without CLEAR pre-processing, emphasizing the utility of this step for extraction of reliable data in RNAseq analysis (**Figure 4E**). Comparison of DEGs identified in the CLEAR processed 10 pg, 100 pg, and 1 ng RNAseq datasets revealed 41.5% of DEGs overlapped between the 100 pg and 1 ng datasets, while only 9.0% of the 10 pg and 1 ng DEGs overlapped. 100 pg and 10 pg shared only 10.6% of DEGs (**Figure 4F**). Analysis of DEG gene counts as percentiles relative to average transcript counts for all genes further revealed that the 100pg RNA input DEGs had a median transcript count level comparable to that of the 1ng input while the 10pg RNA input dataset exhibited a significantly higher percentile of gene expression level (**Figure 4G**). Cumulatively, these data indicate that using 100pg of input RNA preserves data quality and many DEG characteristics of sequencing at the 1ng+ level without requiring its substantially higher number of cells. 10 pg of RNA, in contrast, shows greater variability across biological replicates and has less breadth in the count level of detected DEGs.

### RNAseq enables DEG discovery from FACS isolated NSCs and IPCs from individual mouse hippocampi

Standard bulk sequencing of adult hippocampal NSCs, a particularly sparse in vivo population, requires pooling of several mice to generate sufficient quantities of input RNA. Our findings suggest that when using CLEAR pre-processing, RNA input can be decreased substantially in an lcRNAseq approach and still yield reliable DEGs from a wide range of gene count levels. To test whether an RNAseq approach would be useful for transcriptional sequencing of in vivo NSCs, we used FACS to isolate NSCs and IPCs from 3 individual NestinGFP transgenic mice (Mignone et al. 2004) (**Figure 5A**). Using immunofluorescence of fixed tissue sections of adult NestinGFP mice, we confirmed that GFAP+SOX2+ radial glia like (RGL) NSCs and SOX2+ IPCs, but not DCX+ neuroblasts/immature neurons or NEUN+ mature neurons, expressed NestinGFP (**Supplementary Figure 3A–D**). We also confirmed NestinGFP expression in CD31+ endothelia and OLIG2+ oligodendroglial cells, as expected based on previous work (**Supplementary Figure 3E–F**) (Artegiani et al., 2017). To exclude the endothelia and oligodendroglial cells, we selected for cells immunonegative for CD31, O1 and O4. To specifically capture NSCs and IPCs separately, we used EAAT1 immunolabeling, with NestinGFP+EAAT1+ cells representing NSCs and NestinGFP+EAAT1− cells representing IPCs. To maximize RNA integrity, cells were sorted directly into lysis buffer and converted to cDNA libraries without an intervening RNA isolation step. Direct cDNA synthesis prevented measurement of RNA yield to compare with our in vitro studies, but we used the equivalent of a 60 cell RNA input amount (estimated to approximate 100-200pg input) to generate 3 technical replicates for each biological (mouse) replicate with a 300 cell complexity level for RNAseq and CLEAR filtering (**Figure 5A**). For comparison, we also sequenced in parallel 1 ng of RNA isolated from whole DG of 3 separate mice.

**FIGURE 5.**
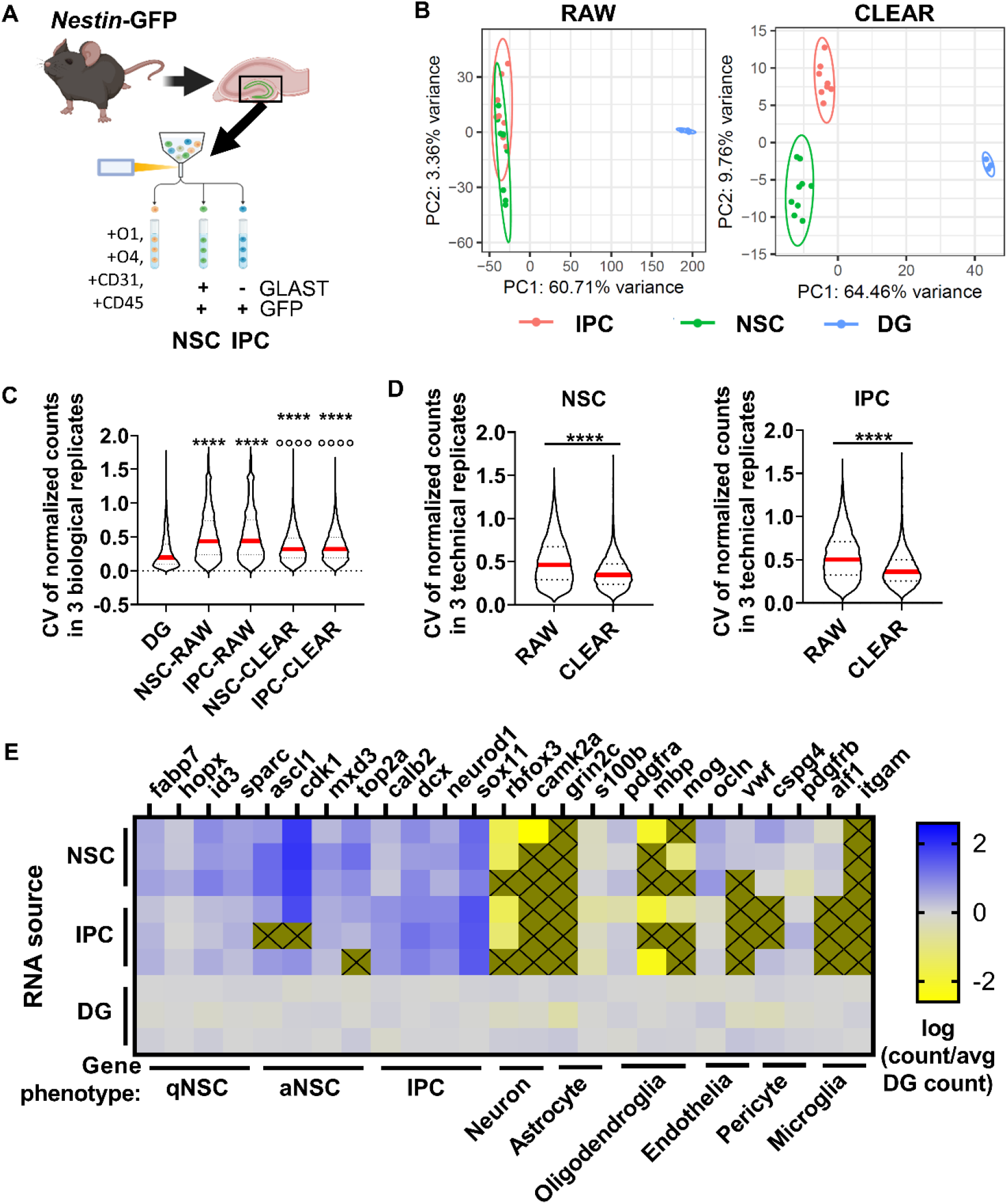
DEG identification in vivo. **(A)** Workflow for DEG identification from hippocampal NSCs and IPCs of individual adult mice. DGs from Nestin-GFP+ mice were isolated and homogenized into single cell suspensions enabling FACS isolation of NSCs and IPCs for RNAseq. **(B)** PCA of transcriptome using unfiltered RAW and CLEAR filtered data. CLEAR filtering improves separation and variance between NSC and IPC samples. **(C)** CV for NSC and IPC biological replicates before and after CLEAR filtering. Dunn’s multiple comparisons ****p<0.0001 vs DG °°°° p<0.0001 vs raw RGL and raw IPC. **(D)** CV for NSC and IPC technical replicates (3/mouse) before and after CLEAR filtering. Unpaired t-test ****p<0.0001. **(E)** Heatmap shows expression of phenotypic genes in each of 3 biological replicates for NSCs, IPCs and whole DG, which was sequenced from 3 separate Wt mice in parallel. High expression of RGL phenotypic markers in NSCs and high expression of IPC phenotypic markers in IPCs confirmed cell type identities. Low expression of neuronal, astrocytic, oligodendroglial, endothelial, pericyte and microglial phenotypic genes confirmed exclusion of other major DG cell types. Gold boxes with an X were not detected.

Principal component analysis of NSCs, IPCs, and whole DG revealed that CLEAR pre-processing both decreased the percent variance within NSCs and IPCs compared to PCA performed on raw data, and also separated NSCs from IPCs into non-overlapping populations (**Figure 5B**). Unfiltered (RAW) CVs for NSCs and IPCs across biological replicates were also more than double the whole DG CV (**Figure 5C**). CLEAR filtering reduced CVs by over 25% for NSCs and IPCs. CLEAR filtering also reduced the CV between technical replicates within the NSCs (by 25%) and IPCs (by 28%) (**Figure 5D**). Because tissue processing and handling can introduce variability when assessing freshly isolated cells, it was not surprising that the technical RNA input in vitro. However, the significant improvement in CVs after CLEAR application confirm its utility in improving transcriptional data from limited starting material.

FACS-isolated NSC and IPC population identities were confirmed with expression of characteristic cell type markers for NSCs and IPCs (**Figure 5E**). Higher expression of quiescent radial glial-like cell (qRGL) markers *Vim* and *Id3* as well as activated radial glial-like cell (aRGL) markers *Neurog2* and *Ascl1* were observed in the NSC samples compared to the IPC and whole DG samples, confirming accurate FACS isolation of NSCs (**Figure 5E**). Likewise, higher expression of intermediate progenitor cell (IPC) markers such as *Calb2* and *Sox11* in the IPC samples compared to the NSC and whole DG samples confirm accurate FACS isolation of IPCs (**Figure 5E**). Exclusion of other cell populations was confirmed via expression of phenotypic genes for neurons, astrocytes, oligodendroglia, endothelia, pericytes and microglia (**Figure 5E**). 177 DEGs were identified between NSCs and IPCs acutely isolated from adult mouse DGs (**Supplementary Table 3**). All together, these findings indicate that RNAseq of adult DG NSCs and IPCs can be achieved from a single mouse per sample replicate with data quality similar to that derived from more bulk-like RNA sequencing.

**Supplementary Figure 3.**
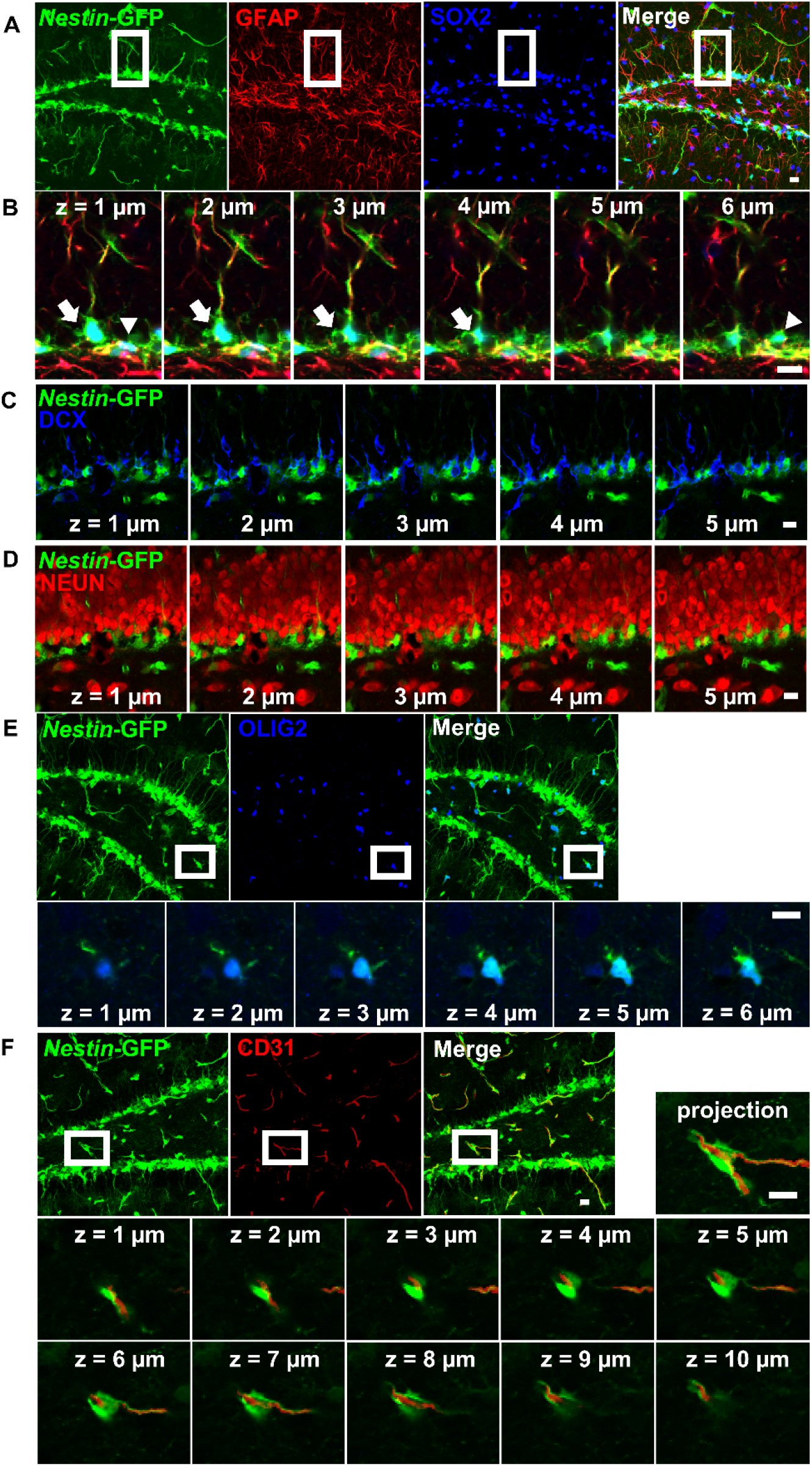
Immunostaining adult Nestin-GFP+ mouse DG demonstrating in vivo DG cell types. **(A)** RGL NSCs are Nestin-GFP+, GFAP+, SOX2+ while IPCs are Nestin-GFP+ and SOX2+ but GFAP-**(B)** Serial z-stack images show the radial glial like process extending from an NSC (white arrow) in the subgranular cell layer through the granule cell layer of the DG. IPCs(arrowhead) are do not have a radial glial like process. SOX2+GFAP+stellate morphology astrocytes were not observed to co-label with Nestin-GFP. **(C)** DCX+ immature neurons are not Nestin-GFP+. **(D)** NEUN+ mature neurons are similary Nestin-GFP−. **(E)** Olig2+ oligodendroglia were co-labeled for Nestin-GFP. **(F)** CD31+ endothelial cells also co-labeled for Nestin-GFP.

### RNAseq identifies DEGs in NSCs and IPCs induced by lateral fluid percussive brain injury

As proof of principle, we applied our RNAseq workflow to a LFPI model of TBI and identified DEGs in adult mouse NSCs and IPCs in vivo (**Figure 6A**). NSCs and IPCs from the DGs ipsilateral to injury of 3 individual mice were FACS isolated 4 hours after LFPI or sham treatment. Whole DGs from separate, but similarly treated, mice were also processed for comparison by 1 ng RNA input sequencing. A total of 6319 genes were identified in NSCs, IPCs, and whole DGs (**Supplementary Table 4**). 23 DEGs were identified in NSCs after LFPI with 15 significantly upregulated and 8 significantly downregulated (**Figure 6B**). In the IPC population, 5 genes were significantly upregulated while 13 genes were downregulated in LFPI mice compared to sham mice (**Figure 6C**). In whole DG, 188 DEGs were identified with 106 significantly upregulated and 82 significantly downregulated in LFPI mice compared to sham mice (**Figure 6D**). Comparison of the DEGs identified in NSCs and IPCs following LFPI did not overlap with DEGs identified on a whole DG level and NSC and IPC DEGs only shared one gene in common (*Slc5a3*), emphasizing the importance of examining individual cell types, even for cells as closely related as NSCs and their IPC progeny. RNAscope fluorescent in situ hybridization (FISH) combined with immunohistochemical staining verified transcriptional upregulation of *Slc5a3* in NSCs and showed a similar trend in upregulation in IPCs following LFPI (**Figure 6E and F**). We also verified two other upregulated DEGs using RNAscope FISH: *Serpina3n* (**Figure 6G**) and *Timp1* both trended higher in expression in NSCs of LFPI mice than sham 4h after injury. (**Figure 6H**). These findings indicate that RNAseq of acutely isolated, in vivo NSCs and IPCs can reliably identify DEGs in an injury model.

**FIGURE 6.**
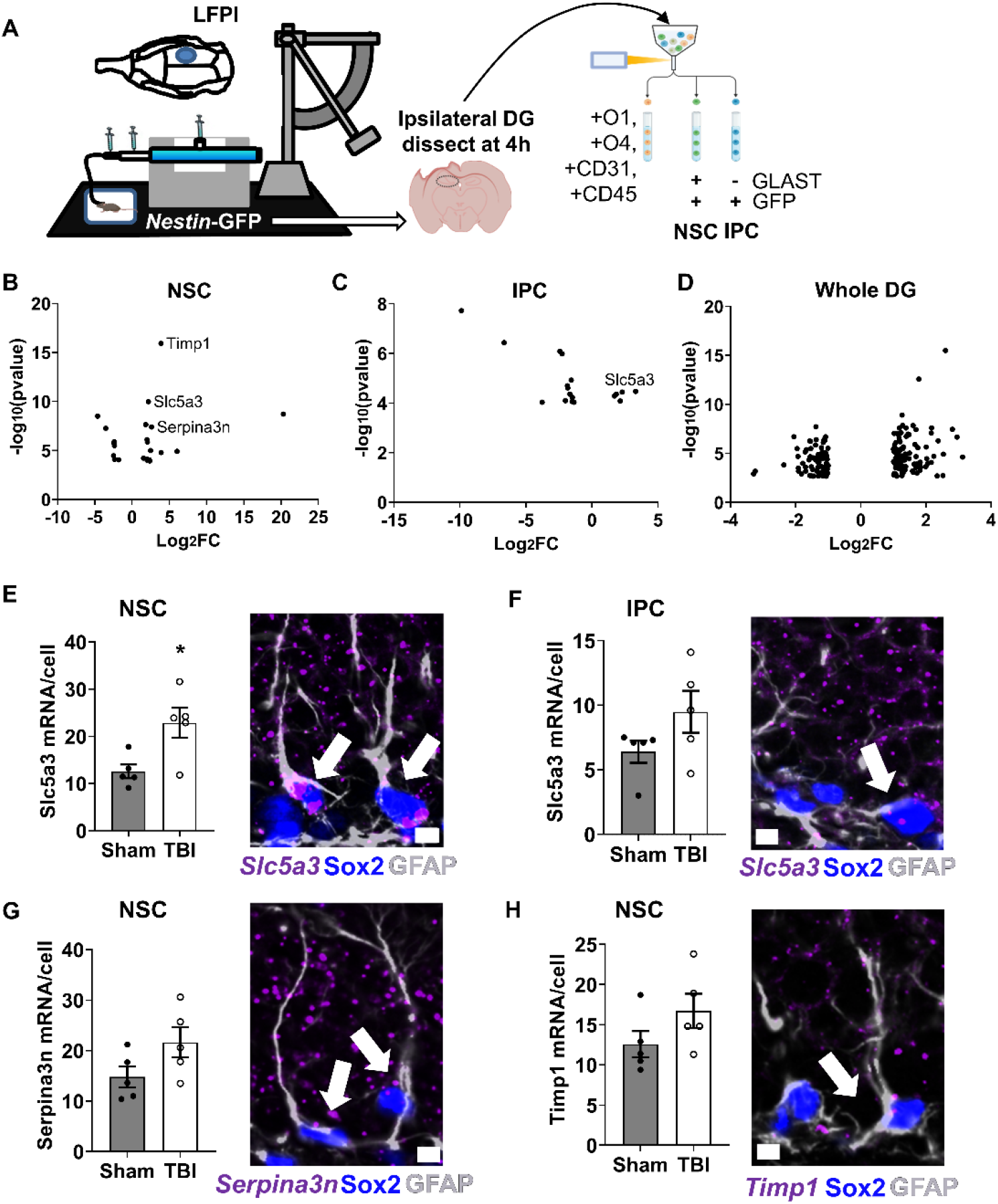
RNAseq and CLEAR filtering enable accurate DEG identification in NSCs and IPCs in mice with lateral fluid percussive brain injury (LEPI). **(A)** Mice (n=3 per group) were given lateral fuid percussive injuries or sham treatment. Four hours after injury, DGs from the ipsilateral side of the brain were isolated and processed for RNAseq as described in Figure 5A. **(B)** Volcano plot of DEGS identified in NSCs after LFPI. **(C)** Volcano plot of OEGs identified in IPCs after LFPI. **(D)** Volcano plot of DEGS identified in whole DG after LFPI. **(E)** Quantification of *SicSa3* mRNA via RNAscope in NSCS (identified via GFAP+SOX2+ immunolabeling) confirmed upregulation after LFPI (lef). p=0.0185, Mean ± SEM of mean RNA puncta per NSC area is shown with 5 individual mice replicates. Representative image with arrows pointing to NSCs with *Sic5a3* expression (right). **(F)** Quantification of *Sle5a3* mRNA in IPCs similarly showed a trend in increased expression after LFPI (lef). p=0.1324. Representative image with arrow pointing to IPC with *Slc5a3* expression (ight). **(G)** Quantification of *Serpina3n* mRNA in NSCs show a trend for upregulation after LFPI (let).p=0.0954. Representative image with arrow pointing to NSC with *Serpina3n* expression (ight). **(H)** Quantification of *Timp1* mRNA in NSCs showed increased expression that did not reach significance (p=0.1627) after LEPI (et). Representative image with arrow pointing to NSC with *Timp1* expression (right). (E-H) n=5 per group, unpaired t-test, *p<0.05. Scale bars = 5 μm.

## Discussion

Endogenous adult hippocampal NSCs provide a source of both cellular and biochemical support for tissue homeostasis. Characterizing these cells at baseline, as well as after injury, may lead to therapeutically relevant strategies for promoting optimal brain function. However, studying adult DG NSCs is challenging due to their relatively low cell number, residence within a complex niche, and their inherent heterogeneity. To overcome this challenge, researchers have increasingly turned to scRNAseq approaches for transcriptional profiling (Hochgerner et al., 2018; Knight and Serrano, 2018; Kulkarni et al., 2019; Shin et al., 2015; Zeng et al., 2018). However, we show here that, despite the many strengths of this approach, scRNAseq may not be the ideal method to answer certain types of research questions. Specifically, we show that scRNAseq analysis of differential gene expression may miss genes that are more moderately expressed and show large fold changes in expression. We show that an lcRNAseq approach can be used to help circumvent this problem and discover DEGs from a broader count range and with greater fold changes, and that such an approach can be adapted for transcriptional profiling and DEG identification in NSCs acutely isolated from adult mouse DG.

Using a model of oxidative stress in cultured NSCs, we identified DEGs from the same source samples using both an scRNAseq and 1ng RNAseq approach. By using cells from the same biological replicates for both sequencing platforms, we avoided potential variation in gene expression induced by a difference in tissue processing. Yet, we found little overlap in the identification of DEGs by these two methods. Both methods yielded accurate DEGs that could be confirmed with qRT-PCR. Furthermore, GO analysis of the DEGs identified using the two different approaches implicated mostly the same biological pathways being triggered by injury. However, the individual genes identified by each method were mostly unique within each platform. This lack of overlap was maintained when scRNAseq data were analysed without filtering for high or low feature counts and when the same statistical method was used for DEG discovery in both datasets, suggesting it was not an artefact of data processing/analysis.

When we probed the difference in the kinds of DEGs identified by scRNAseq versus 1 ng RNAseq, we found that scRNAseq identified high count genes with lower fold changes while 1 ng RNAseq identified genes with a wider range of count levels, including many genes with moderate counts and higher fold changes. Two independent scRNAseq datasets from different tissues and experimental designs showed a notably similar DEG profile in terms of relative count level and fold change as in our data. This similarity across such different experiments suggests that the restriction of DEGs to high count genes with lower fold change may be an inherent limitation of scRNAseq via 10x Chromium approach. Importantly, this bias in count level and fold change emerged in comparison to transcripts detected above 0 in each respective dataset. scRNAseq, of course, yields fewer detected genes than a bulk approach but this difference in DEG profile emerged even among genes that would be recorded in the dataset as detected and therefore having been evaluated for potential to be a DEG.

There are several likely contributors to the selectivity of scRNAseq data for generating DEGs from high count transcripts. The first and most obvious potential contributor is the high zero-count rate in scRNAseq data, due to the very low RNA input from a single cell. Transcripts with counts near threshold of detection will be characterized by many 0-count genes and therefore have relatively high variability from cell to cell. This high variability may make such genes less likely to be detected as significantly different between two conditions. Discovery of genes with higher counts, and therefore lower zero-influenced variability, would be favored. By being higher count, fold change is similarly likely to be more constrained by a ceiling effect, yielding lower fold change DEGs. Analytical processing compensations for heteroskedasticity may also play a role. Heteroskedasticity is the phenomenon whereby genes with relatively low counts exhibit higher fold changes. Thus, general DEG analysis methods, such as DESeq2, and scRNAseq analysis pipelines, such as Seurat, correct for this problem by applying a variance-stabilizing preprocessing step that transforms the data and minimizes the effect of count-based technical noise on ratio-based outputs such as log fold change in gene expression (Ahlmann-Eltze and Huber, 2021; Love et al., 2014). Although heteroskedasticity is recognized for any count-based data, standard bulk level RNAseq is inherently more robust and delivers generalized expression data for hundreds of thousands of cells. This facilitates accurate correction for heteroskedasticity, while in scRNAseq, low biological replicates and very low input RNA amounts may lead to over-correction of true variation in gene expression by variance-stabilizing preprocessing (Love et al., 2014; Mou et al., 2020).

While our findings suggest that scRNAseq and low-input population level RNAseq are both similarly accurate, as measured by replicability, they suggest different utility of each approach depending on experimental goal. If characterization of cellular heterogeneity is the goal, scRNAseq is obviously superior. However, discovery of genes that are moderate to lowly expressed in cells yet exhibit higher fold changes upon stimulation is also a common goal, particularly when searching for candidate molecular mediators or druggable targets in models of injury and disease. Relying on scRNAseq alone for DEG discovery could therefore limit the scope of understanding of disease or injury-associated transcriptional signatures. The difference in DEG characteristics between scRNAseq and 1ng RNAseq shown here emphasizes the importance of identifying the ultimate goals and readouts of an experiment to choose the method that best addresses the needs of the study.

Although several studies have provided invaluable insight into hippocampal NSC biology using scRNAseq (Artegiani et al., 2017; Hochgerner et al., 2018; Shin et al., 2015), standard bulk RNAseq to study endogenous adult hippocampal NSCs has been challenging, as evidenced by the dearth of such studies. Though NSCs have been successfully profiled when combined with their IPC progeny via bulk RNAseq approach (Adusumilli et al., 2021), NSCs alone are sufficiently rare that their “bulk” level transcriptional profiling is especially difficult. Here, we optimized a protocol that enables population level transcriptional analysis of this rare cell type from individual mice. First, we used cultured adult hippocampal NSCs to determine 100 pg as a lower RNA input amount that enables profiling of DEGs that are comparable to those obtained with standard bulk sequencing. Using this threshold of 100 pg RNA input as a guide for in vivo analysis, we isolated adult DG NSCs and IPCs from individual Nestin-GFP reporter mice to profile the transcriptomes of each population at a 300 cell complexity level.

To ensure accurate identification of DEGs from this low input level, we applied CLEAR filtering which was previously shown to minimize technical noise due to limited RNA input (Walker et al., 2020). In brief, CLEAR preprocessing removes transcripts from analysis that are detected below a threshold. That threshold is determined by observing where on an mRNA transcript sequence RNAseq read fragments map. When reads show preferential mapping to only 3’ and 5’ ends of their mRNA transcripts, it indicates strong RNA degradation, a feature which predominates as transcript count drops. CLEAR filtering removes transcripts with counts below the threshold where most transcripts start to show this pattern of mapping more exclusively towards the 3’ and 5’ ends of their mRNA sequences. CLEAR filtering of our data improved the coefficient of variation between biological replicates for normalized counts at all RNA input amounts in vitro and in vivo, and was essential for effective transcriptional separation of in vivo-isolated NSCs and IPCs via PCA. We showed that this workflow, with CLEAR filtering, accurately profiled the transcriptomes of NSCs and IPCs. This ability to capture population level complexity with a low amount of RNA input is valuable when studying rare cell populations in complex experimental models of disease or injury where maximizing biological replicates is critical but limited by time and labor costs.

To demonstrate the utility of our approach for transcriptional profiling of in vivo NSCs, we applied it in a proof-of-principle experiment identifying TBI-associated DEGs from in vivo NSCs and IPCs. Using our workflow, we obtained cell type specific DEGs four hours after the LFPI model of TBI. In response to LFPI, NSCs and IPCs showed mostly unique DEGs, with only one exception: both upregulated *Slc5a3*, a sodium-coupled inositol transporter protein that maintains osmotic pressure in cells and regulates intracellular myo-inositol levels (Andronic et al., 2015). We confirmed transcriptional upregulation of *Slc5a3*, as well as two other DEGs, *Serpina3n* and *Timp1*, with RNAScope in situ hybridization paired with immunofluorescent staining to identify NSCs and IPCs. These findings suggest that our population-level approach to RNAseq of isolated NSCs and IPCs can effectively identify replicable changes in gene transcription after an injury stimulus.

There are several limitations to this study. First, we only compared bulk RNAseq to scRNAseq using a 10x Chromium platform. It is therefore possible that other approaches to scRNAseq would yield different results than what we found. However, many of the limitations we noted seemed inherent to working with very low RNA input levels and were not exclusive to our experimental platform (in vitro NSCs) or any one approach to data analysis. Second, the specific cell/RNA input levels that we identified as yielding more bulk-like range in DEG count level and fold change may not apply outside of our selected cell population (adult NSCs). RNA content, cellular heterogeneity and capture efficacy of different cell types and tissue sources will likely influence the appropriate cell input needed in other models. Third, transcriptomics does not equate with proteomics. Particularly in NSCs, there appears to be substantial translational priming, in which mRNA is produced but not translated (Denninger et al., 2020; Kjell et al., 2020). Thus, both transcriptional and proteomic analyses are needed to accurately characterize NSCs in health and disease.

In conclusion, we present a comparison of two different approaches to transcriptional profiling, scRNAseq and population level RNAseq, of adult hippocampal NSCs, a rare cell type that is difficult to study in vivo. We found that each had their strengths, as well as weaknesses, which should be balanced with the needs of each specific study. We have shown here that our method for in vivo transcriptional profiling in a bulk-like lcRNAseq approach can provide valuable information about rare cell populations that are traditionally difficult to study in vivo. Thus, we present our workflow as an addition to the transcriptional toolbox for studying limited in vivo cell types moving forward.

## Materials and Methods

### Animals

NestinGFP mice (Mignone et al., 2004) (Jackson Labs, #033927) and Wt C57Bl/6J mice (Jackson Labs, #000664) were housed in a 12 hour light-dark cycle with food and water ad libitum. To isolate brains for fluorescence in situ hybridization, adult mice (6-9 wk old) were anesthetized with an intraperitoneal injection of ketamine (87.5 mg/kg) and xylazine (12.5 mg/kg). Mice were then transcardially perfused with ice cold PBS followed by cold 4% PFA. For immunofluorescence and whole DG RNA isolation, mice were only perfused with ice cold PBS. For lcRNAseq experiments, mice were only perfused with ice cold HBSS without calcium or magnesium. This study was approved by the Institutional Animal Care and Use Committee (IACUC) at the Ohio State University in accordance with institutional and national guidelines.

### DG isolation and FACS for RNAseq

To isolate DGs for subsequent fluorescence activated cell sorting (FACS), adult mice (6-9 wk old) were anesthetized and perfused with HBSS as described above. Following perfusion, brains were removed and placed in cold HBSS on ice. To expose the hippocampus, brains were bisected along the midsagittal line and the cerebellum and diencephalic structures were removed. Under a dissection microscope (Zeiss), the DG was excised using a beveled syringe needle and placed in ice cold HBSS without calcium or magnesium. DGs were then mechanically dissociated with sterile scalpel blades before enzymatic dissociation with a pre-warmed papain (Roche 10108014001)/dispase (Stem Cell Technologies 07913)/DNase (Stem Cell Technologies NC9007308) (PDD) cocktail at 37°C for 20 min. Afterwards, the tissue was again mechanically disrupted by trituration for 1 min. Dissociated cells were collected by centrifugation at 500g for 5 min before resuspending in HBSS without calcium/magnesium. Cells were then filtered through a 35μm nylon filter before staining with fluorescent antibodies (**Supplementary Table 5**) on ice for 30 minutes. During the last 10 minutes of staining, Hoechst dye was added for live/dead discrimination. All cells were washed twice following staining and immediately sorted as NSC or IPC populations based on fluorescent markers with the FACSAria III (BD Biosciences). CD31-, CD45-, O1-, and O4 negative live cells were designated as NSCs if double positive for GLAST and GFP or IPCs if GFP positive and GLAST negative. Three technical replicates of 300 cells each were sorted from each individual mouse into 1.5mL microcentrifuge tubes containing cell lysis buffer buffer from the Clontech SMART-Seq HT (Takara) kit for direct cDNA synthesis and RNAseq library generation.

### Cell culture

NSCs were isolated from adult DGs of C57Bl6/J mice as described in (Babu et al. 2011). Two separate lines, one from 4 pooled C57Bl6/J male mice and one from 4 pooled C57Bl6/J female mice, were used in experiments between passage 5 and 15. NSCs were cultured on poly-D-lysine (Sigma) and laminin (Invitrogen) coated plates in Neurobasal A media (Invitrogen) with 1x B27 supplement without vitamin A (Gibco), 1x glutamax (Invitrogen) and 20 ng/ml each of EGF and FGF2 (Peprotech). There were no inherent differences in morphology or proliferation between NSC cultures and both lines differentiated into neurons and glia upon culture in differentiation conditions, as we previously showed (Denninger et al., 2020). All cultures were verified to be mycoplasma-free. For oxidative stress experiments, NSCs in triplicate were treated at 70% confluency with 500 μM H_2_O_2_ (Sigma) or equal volume of vehicle (PBS) for 24 hours. All cells were harvested with brief accutase treatment and one wash with HBSS for RNA isolation or scRNAseq.

### RNAseq of cultured cells and whole DG

RNA from 30,000 cultured adult NSCs or whole DGs were isolated with the Clontech Nucleospin RNA XS Plus isolation kit (Takara 740990.10) per manufacturer protocol. RNA quality (RNA Integrity Number or RIN) and quantity was assessed using Agilent BioAnalyzer RNA 6000 Pico Kit and the Invitrogen Qubit RNA HS Assay kit (Invitrogen), respectively. All cultured samples used a RIN value of 10 while whole DG samples had RIN values over 8. RNA from cultured NSCs was serially diluted to 10-, 100-, and 1000-pg for RNA input quantity studies. Whole DG libraries for bulk RNAseq were generated with the NEBNext Ultra II Directional RNA Library prep kit (New England Biolabs). The Clontech SMART-Seq HT (Takara) kit was used for global preamplification of cultured and 20% (~60 cell input) of the 300 FACS-isolated NSC cell lysate for low input RNAseq. Purified library products were then used in HiSeq 4000 paired-end sequencing (Illumina) to a depth of 15–20 million 2 × 150 bp clusters. FASTQ files generated for each library were trimmed using AdapterRemoval v2.2.0 (Schubert et al., 2016), ensuring that all sequencing adapters were removed and that the average quality score for each read was above Q20 (representing 1 in 100 Illumina base error rate). Reads which were aligned by HISAT2 v2.0.6 (Kim et al., 2015) against rRNA, mtDNA, or PhiX bacteriophage (Illumina spike-in control) sequences, retrieved from NCBI RefSeq (O’Leary et al., 2016), were removed from each FASTQ file, as these do not represent gene expression signal. All remaining reads were aligned against the reference mouse genome GRCm38p4 with HISAT2. The resulting BAM alignment files were sorted and indexed before further analysis.

Alignments were quantified using the featureCounts utility from the Subread package v1.5.1 (Liao et al., 2014, 2013) in unstranded mode using GENCODE (Harrow et al., 2012, 2006) mouse gene reference version M14 in GTF format. Custom Python scripts were used to produce a formatted gene expression counts table from the raw output of featureCounts. RNAseq Quality metrics were derived using a modification of the QuaCRS quality control workflow (Kroll et al., 2014) which includes running RNA-SeQC v1.1.8.1 (DeLuca et al., 2012), FASTQC v0.11.5, and RSeQC v2.6.2 (Wang et al., 2012). Finally, coverage maps of each BAM file were derived using the Bedtools ‘genomecov’ utility v2.27.0 (Quinlan and Hall, 2010).

RNAseq coverage maps were processed with the CLEAR v1.0 (Walker et al., 2020) workflow to determine which genes were reliably quantified. In brief, for each transcript in the UCSC GRCm38 release (Kuhn et al., 2013), a parameter (μi) is calculated, which represents the positional mean of the reads covering the transcript, normalized to the length of each sequence. Sequential bins of 250 μi values each, ordered by descending expression, are fit to a sum of two beta distributions (Gupta and Nadarajah, 2004) for the determination of two free parameters, which are thresholded to determine genes which “pass” CLEAR. Unless otherwise noted, these passing genes are used for downstream analysis. DEGs were derived from filtered RNAseq expression counts tables following the DESeq2 v1.20.0 protocol (Love et al., 2014) implemented in R v3.5.0. DEGs were then processed with the pcaExplorer v2.6.0 (Marini and Binder, 2019) visualization package to produce principal component analysis (PCA) projections using the default settings on r-log transformed counts.

### scRNAseq

30,000 NSCs were pooled from triplicate H_2_O_2_− or vehicle-treated cultures and loaded onto the 10X Genomics single cell sequencing platform using the standard kit. The 3’ RNA-seq library was sequenced using paired-end 150bp approach on an Illumina HiSeq 4000 sequencer. Data from vehicle-treated cells was previously published in (Denninger et al., 2020) and similar analysis was performed here, but on the combined data, including both vehicle-treated and H_2_O_2_-treated cells. CellRanger v3.0.2 (Zheng et al., 2017) was used to demultiplex, align, and deduplicate sequencing reads in BCL files. Single-cell data in feature-barcode matrices were then processed using Seurat v3.0.1’s default pipeline (Butler et al., 2018; Stuart et al., 2019) to identify unsupervised cell clusters and generate a uniform manifold approximation and projection (UMAP) plot. Seurat’s feature expression analysis was used to visualize genes markers known to be expressed in cell populations displayed in the UMAP plot.

### qRT-PCR

Cultured NSCs were lysed in culture plates and RNA was isolated with the BioRad Aurum™ Total RNA Mini Kit according to the manufacturer protocol. Isolated RNA was quantified and assessed for quality using the BioTek Epoch Microplate Spectrophotometer. cDNA was synthesized with the BioRad iScript™ cDNA Synthesis Kit in the ThermoFisher Applied Biosystems 2720 Thermal Cycler according to manufacturer protocol. qRT-PCR was performed in the BioRad CFX96 Touch Real-Time PCR Detection System with BioRad SsoAdvanced Universal SYBR Green Supermix and primers listed in Supplementary Table 5. ΔΔCt values generated with normalization to housekeeping gene Rpl7 then converted to fold change (relative to vehicle).

### Lateral fluid percussion injury

All surgical procedures were performed as previously described (Tapp et al., 2020). Briefly, 6–9-week-old mice were anesthetized with 4% isoflurane gas in an induction chamber for 4 min. Mice were then positioned in a stereotaxic frame before making a sagittal incision to expose the cranium. Midway between bregma and lambda on the right parietal bone, a 3.0-mm craniectomy was trephined, leaving the intact dura mater exposed. A modified portion of a Leur-Loc syringe (3.0-mm inside diameter) was secured over the craniectomy site with cyanoacrylate adhesive. Mice were placed in their home cages on a heating pad to recover. Once mice resumed normal activity, mice were returned to the vivarium for 24 hours. The next day, mice were anesthetized with 4% isoflurane in an induction chamber for 4 min. Using the modified Leur-Loc syringe, mice were connected to the fluid percussion injury device (Custom Design & Fabrication). For mice designated to the TBI group, a prepositioned pendulum was released onto the end of the LFPI device to deliver a fluid pulse onto the exposed dura mater, inducing a moderate LFPI. Sham mice were attached to the LFPI device but did not receive a fluid pulse. The modified syringe and adhesive were removed following LFPI or sham treatment. The incision was stapled closed. All animals were placed on a heating pad and injury severity was assessed with the self-righting reflex test. After the subjects demonstrated the righting reflex, they were returned to their home cages on a heating pad. 4 hours after injury, mice were perfused for either histology, RNA isolation, or FACS.

### Histology (IF and RNAscope)

For identification of in vivo cell types, brains from NestinGFP mice were harvested and fixed overnight at 4°C in 4% PFA before overnight equilibration in 30% sucrose. Serial 40μm sections were rinsed in PBS before blocking with 1% normal donkey serum (Jackson) and 0.3% Triton X-100 in PBS for 30 minutes at room temperature. Sections were incubated with primary antibodies (**Supplementary Table 5**) overnight at 4°C in blocking solution on an orbital shaker. The next day, sections were washed with PBS and incubated with fluorescently conjugated secondary antibodies diluted in blocking solution (**Supplemental Table 5**) for 2h at room temperature and counterstained with Hoechst 33342 (1:2000) for nuclear visualization. Sections were then washed and mounted onto slides before coverslipping with ProLong Gold anti-fade solution (Molecular Probes). Slides were imaged in 1-μm z-stacks on an LSM700 confocal microscope (Zeiss) with a 40× oil objective.

For RNAscope, four hours after LFPI or sham injury, brains from NestinGFP mice were harvested and fixed overnight at 4°C in 4% PFA before serial overnight equilibration in 10%, 20%, and 30% sucrose. Fixed equilibrated tissue was snap frozen in OCT in a dry ice/100% ethanol bath and stored at −70°C. 12 μm cryosections, 1 section per slide, were prepared with a cryostat. Slides were stored at −70°C with desiccant until staining. RNA in situ hybridization was performed with RNAscope Multiplex Fluorescent v2 Assay (Advanced Cell Diagnostics) according to manufacturer recommendations for using fixed frozen tissue samples with the following modifications to enable concurrent immunohistochemical staining. The pretreatment steps were replaced with a 15 min modified citrate buffer (Dako) antigen retrieval step in a steamer at 95°C. To enable subsequent immunohistochemical staining, the protease III step was excluded. Probes for mouse Slc5a3 (ACD custom design NPR-0006102), mouse Serpina3n (ACD 430191), and mouse Timp1 (ACD 316841) RNA were hybridized to tissue prior to immunohistochemical staining for GFAP and SOX2 protein. Immunostaining for GFAP and SOX2 was conducted as described above with the following exceptions. Blocking was performed with 10% normal donkey serum in TBS-1% BSA. Antibody incubations were performed in TBS-1% BSA. All washes were performed with TBST. DAPI provided by the RNAscope Multiplex Fluorescent kit was used for nuclear counterstaining. Images were acquired with a Zeiss Axio Observer Z1 microscope with Apotome for optical sectioning using a 20x air objective. Full z-stacks were acquired for analysis. NSCs were identified based on SOX2 positivity and GFAP+ apical processes extending from the nucleus in 1 μm z-stack images from n = 4 mice. IPCs were similarly identified based on SOX2 positivity but without GFAP+ apical processes. mRNA puncta for all 3 genes were counted manually throughout the depth of cell nuclei and length of processes.

### Statistics

Statistics were performed as described in each figure legend. χ2 contingency test was used to compare reproducibility of DEGs identified in cultured NSCs after oxidative stress by scRNAseq and 1 ng RNAseq. For the qRT-PCR confirmation of top DEGs by platform, a Mann-Whitney test was performed to identify individual genes with significant upregulation. To compare percentiles of gene expression by DEGs identified via scRNAseq versus 1 ng RNAseq, all genes with no expression in any cell were first eliminated. Averaged transcript counts for DEGs were compared to averaged transcript counts of all detected genes within each sequencing platform. Unpaired t-test was used to compare scRNAseq and 1ng RNAseq of our cultured NSCs. Dunnett’s multiple comparison test was used to compare our data with data published in [Kang et al. 2018] and [Jin et al. 2021]. To compare coefficient of variation between samples with or without CLEAR filtering, a one-way ANOVA with the Kruskal-Wallis test followed by Dunn’s multiple comparisons or unpaired t-tests were applied. NSC and IPC mean expression per cell of Slc5a3, Serpina3n, and Timp1 were compared using unpaired t-tests. All analyses were performed using Prism (v9.0; GraphPad Software) and p<0.05 was considered significant.

## Author Contributions

JKD: Execution of research and writing. JKD and EDK: Primary contributors to research design, analysis, and writing. EDK: Funding acquisition and supervision. LAW, XC, AT, AP, RB, PY: Contributed to design, execution, analysis, and writing of methods for RNAseq studies. ZT and OKC: Performed LFPI studies. SS and RR: Provided general support in execution of research.

## Acknowledgments

We thank all of the members of the Kirby lab for excellent discussions during the execution of this project as well as planning and writing of this work. We also thank the OSU Chronic Brain Injury Initiative (CBI), the OSUCCC Genomics Shared Resource, the Kokiko-Cochran lab, and the OSU animal care staff for their support.

## Data Availability Statement

RAW RNA-seq sequencing FASTQ files and processed counts tables will be deposited into the NCBI Gene Expression Omnibus (GEO) upon publication.

